# Genome wide profiling of histone H3 lysine 4 methylation during the Chlamydomonas cell cycle reveals stable and dynamic properties of lysine 4 trimethylation at gene promoters and near ubiquitous lysine 4 monomethylation

**DOI:** 10.1101/2021.09.19.460975

**Authors:** Daniela Strenkert, Asli Yildirim, Juying Yan, Yuko Yoshinaga, Matteo Pellegrini, Ronan C. O’Malley, Sabeeha S. Merchant, James G. Umen

**Author notes:** Corresponding author: James G. Umen.

## Abstract

Chromatin modifications are key epigenetic regulatory features with roles in various cellular events, yet histone mark identification, gene wide distribution and relationship to gene expression remains understudied in green algae. Histone lysine methylation is regarded as an active chromatin mark in many organisms, and is implicated in mediating active euchromatin. We interrogated the genome wide distribution pattern of mono- and trimethylated H3K4 using Chromatin-Immunoprecipitation followed by deep-sequencing (ChIP-Seq) during key phases of the Chlamydomonas cell cycle: early G_1_ phase (ZT1) when cells initiate biomass accumulation, S/M phase (ZT13) when cells are undergoing DNA replication and mitosis, and late G_0_ phase (ZT23) when they are quiescent. Tri-methylated H3K4 was predominantly enriched at TSSs of the majority of protein coding genes (85%). The likelihood of a gene being marked by H3K4me3 correlated with it being transcribed at one or more time points during the cell cycle but not necessarily by continuous active transcription. This finding even applied to early zygotic genes whose expression may be dormant for hundreds or thousands of generations between sexual cycles; but core meiotic genes were completely missing H3K4me3 peaks at their TSS. In addition, bi-directional promoters regulating expression of replication dependent histone genes, had transient H3K4me3 peaks that were present only during S/M phase when their expression peaked. In agreement with biochemical studies, mono-methylated H3K4 was the default state for the vast majority of histones that were outside of TSS and terminator regions of genes. A small fraction of the genome which was depleted of any H3 lysine methylation was enriched for DNA cytosine methylation and the genes within these DNA methylation islands were poorly expressed. Genome wide H3K4me3 ChIP-Seq data will be a valuable resource, facilitating gene structural annotation, as exemplified by our validation of hundreds of long non-coding RNA genes.

## INTRODUCTION

Eukaryotic nuclear DNA is organized into chromatin, enabling condensed packaging of DNA in the nucleus. Chromatin can be subdivided into domains of euchromatin and heterochromatin. These domains are defined by their level of compaction and are usually associated with active gene expression for euchromatin and silent, repeat-rich DNA for heterochromatin. Nucleosomes composed of histone octamers and wrapped with DNA are the fundamental packaging unit for chromatin (1-3). Histones are positively-charged, mainly globular proteins with unstructured N-terminal tails that are enriched in posttranslational modifications (PTMs). The identification of specific types of histone modifications at different loci and chromatin domains --including acetylation, methylation, phosphorylation, sumoylation and ubiquitination--suggested that chromatin contains complex regulatory features in addition to its packing function (4). Accordingly, different histone PTMs have been correlated with the repression or activation of associated gene expression (5, 6). Previous studies have demonstrated that histone lysine methylation deposited by histone methyltransferases and recognized by a variety of chromatin “readers” plays an important role in regulating gene expression in diverse eukaryotic taxa. Histone H3 lysine 4 (H3K4) methylation is one of the most studied histone PTM and H3K4 can have up to three different methylation states, mono-, di- or tri-methylation (7).

Methylation of H3K4 has been extensively studied in the budding yeast *Saccharomyces cerevisiae*. In *S. cerevisiae*, tri-methylation of Lysine 4 at histone H3 is a signature motif at the 5′ end of transcribed open reading frames and marks transcription start sites (TSS) of genes that are poised for transcription (8-11). Furthermore, H3K4me3 was also associated with epigenetic memory (9) and maintaining appropriate regulation of gene expression as cells age (12). It was shown that mono-, di-, and trimethylation of H3K4 is mediated by the Set1 subunit of the COMPASS complex in yeast (13-15). Notably, enrichment of H3K4me3 at the 5’ end of open reading frames (ORFs) seems to be conserved in metazoans. In humans, mouse and fly, H3K4me3 is also enriched at TSS of actively transcribed genes (16-19). By contrast, mono-methylation of H3K4 (H3K4me1) was shown to be enriched at enhancers and might play a role in mediating epigenetic memory (20).

In land plants, such as Arabidopsis, Rice and Maize, tri-methylation at lysine 4 of histone H3 occurs predominantly at gene promoters, while mono-methylation at lysine 4 of histone H3 is distributed within transcribed regions in Arabidopsis (21-25). It was also shown that enrichment of H3K4me1 at gene bodies of Arabidopsis genes highly correlated with DNA methylation (21). While H3K4me3 peaks at TSSs seem to be stable after environmental perturbations in most organisms, enrichment of H3K4me3 at promoter regions displayed dynamic changes in response to dehydration stress in Arabidopsis (26).

*Chlamydomonas reinhardtii* (referred to Chlamydomonas hereafter) is a single-celled, eukaryotic green alga and a reference organism for investigating photosynthesis, the cell cycle, ciliary biogenesis, cellular metabolism and other processes (27). Previous chromatographic and mass spectrometry studies of bulk purified histones revealed that nearly all detectable H3 has lysine methylation in Chlamydomonas with very little unmethylated H3 (28-30). The majority of H3K4 was mono-methylated (H3K4me1) with a few percent having tri-methylation (H3K4me3). Moreover, using top-down mass spectrometry, a strong positive correlation was found between H3K4me3 and high levels of H3 acetylation, while H3K4me1 H3 had much lower acetylation levels (30). Mono- and tri-methylation of lysine 4 at histone H3 have been further studied in Chlamydomonas, but mainly in the context of transgene silencing and at single loci using chromatin immunoprecipitation and quantitative PCR (ChIP-qPCR) (31, 32). H3K4me1 was associated with inactive euchromatin at transgenic loci, while H3K4me3 was enriched at promoters of highly transcribed *RBCS2* and *HSP70A* genes (31, 32).

Although some genes in Chlamydomonas are expressed stably and constitutively, most of the transcriptome is expressed in a periodic manner over the course of a diurnal cycle under phototrophic conditions. Under a typical 12hr light:12hr dark diurnal regime, cell growth (G1 phase) occurs during the light period and cell division (S and M phases or S/M) occurs by multiple fission around the time of the light-to-dark transition and is followed by a quiescent period (G0) before the next light phase (Fig. 1) (33, 34). There are also some genes in Chlamydomonas which are expressed infrequently and facultatively such as those required during gametogenesis, mating and the diploid phase of the life cycle (35-37), or those induced under specific stress conditions (38-41). The relationships between these different gene expression patterns (always on, cycling on and off, and expressed infrequently and/or only under very specific conditions) and chromatin architecture have not been examined previously.

**Figure 1.**
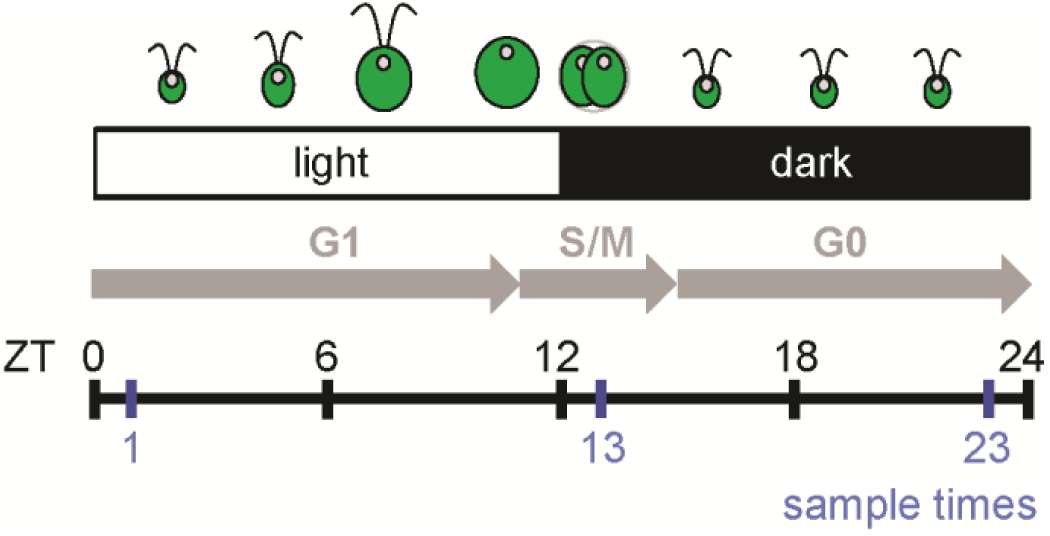
Experimental design and sampling scheme. Diagram shows ZT time of light (ZT0 to ZT12), and dark (ZT12 to ZT24). Shown is a synchronized Chlamydomonas culture with growth (G1), division (S/M), and resting (G0) stages indicated by gray arrows. Cultures were sampled at ZT1 (early G1), ZT13 (mitosis) and ZT23 (end of G0) (blue vertical lines).

Chromatin immunoprecipitation (ChIP) is one of the most widely used tools to discover DNA-protein interactions and when combined with high-throughput sequencing (ChIP-Seq), it enables the discovery of epigenetic regulatory patterns on a genome wide scale. Here we focused on interrogating the methylation status of H3K4 during the vegetative cell cycle using ChIP-Seq under synchronous diurnal conditions. Combining ChIP-seq data for H3K4me3 and H3K4me1 with DNA methylation data and matching transcriptome data allowed us to produce an integrated map of epigenetic dynamics at key cell cycle phases. We used these data to further refine hypotheses about how the epigenetic landscape is defined and remodeled during the Chlamydomonas cell cycle. We further established how promoter enrichment of H3K4me3 can be used as a tool to facilitate annotation of novel protein coding genes as well as long non-coding RNAs (lncRNAs).

## RESULTS

### Comprehensive analysis of Chlamydomonas H3K4me3 and H3K4me1 during the vegetative cell cycle

We investigated two histone modifications (H3K4me1 and H3K4me3) in Chlamydomonas using ChIP-Seq, with input DNA serving as a control, and using antibodies that were previously shown to be effective for ChIP in Chlamydomonas (31). We carried out ChIP-Seq at three key stages of a synchronous diurnal cycle under phototrophic conditions—early G1 phase (ZT1) when cells are actively photosynthesizing and growing in biomass, S/M phase (ZT13) when cells are rapidly dividing, and late G0 phase (ZT23) when cells are undergoing dark phase metabolism and photosynthesis is inactive (Fig 1). Matched RNA-seq data from triplicate samples was obtained from a previous study (33) and analyzed along with ChIP-Seq data from this work. The majority of reads (> 91%) could be mapped to the nuclear genome, which suggested that the sequencing data were of a high quality (Supplemental table 1). Model-based analysis using MACS software (42) was used for peak identification. We identified 13771, 14032 and 13733 H3K4me3 peaks in ZT1, ZT13 and ZT23, respectively (Table 1). Notably, replicate samples from all three time points showed excellent correlation with each other, with the majority of peaks (>98%) being identified in both replications (Table 1). Visual inspection of the data obtained for H3K4me1 on the genome browser revealed that the majority of the genome seems to be mono-methylated at H3K4 with no clear peak enrichment, making traditional peak calling impossible. This near featureless landscape of H3K4me1 was not due to a technical failure as we identified valleys of H3K4me1 depletion that overlapped with H3K4me3 peaks (Table 1).

**Table 1.** Peak calling statistics.

| Time point | Agreement between replicates | Total consensus peaks/depleted regions | Number of peaks w/gene overlap |
| --- | --- | --- | --- |
| H3K4me3, ZT1 | 98.5% | 13771 | 12754 (92.6%) |
| H3K4me3, ZT13 | 99.7% | 14032 | 12903 (92.0%) |
| H3K4me3, ZT23 | 99.0% | 13733 | 12733 (92.7%) |
| H3K4me1, ZT1 | 91.3 % | 10348 | 9980 (96.4%) |
| H3K4me1, ZT13 | 94.0% | 12144 | 11377 (93.7%) |
| H3K4me1, ZT23 | 91.4% | 9523 | 9148 (96.1%) |

### Distribution of H3K4me3 and H3K4me1 in the Chlamydomonas genome

Notably, the majority of H3K4me3 peaks (∼85%) overlapped with the transcription start sites (TSS) and most of the remaining peaks were intergenic (∼11%) or found near 3’ ends of genes (∼3%) (Fig. 2A). The overlap with TSS regions suggested that these were focal points for H3K4me3, a result that was apparent when the relative enrichment for this mark was plotted by genic position across all gene models (Fig. 2B). The general pattern was enrichment at the predicted TSS, modest depletion in the gene body, and a small peak just downstream of the transcription termination site. While H3K4me1 was not enriched in any specific genic region, it was depleted at promoter regions and inversely correlated with H3K4me3 enrichment (Fig. 2C). To gain a better understanding of the gene wide distribution of the two chromatin marks, we plotted the sizes of H3K4me3 peaks individually for all 17741 predicted protein coding genes and ordered them from strongest to weakest. This plot shows that around 90% of predicted genes have a H3K4me3 peak around their TSS with a proportionately deep valley for H3K4me1 (Fig. 3).

**Figure 2.**
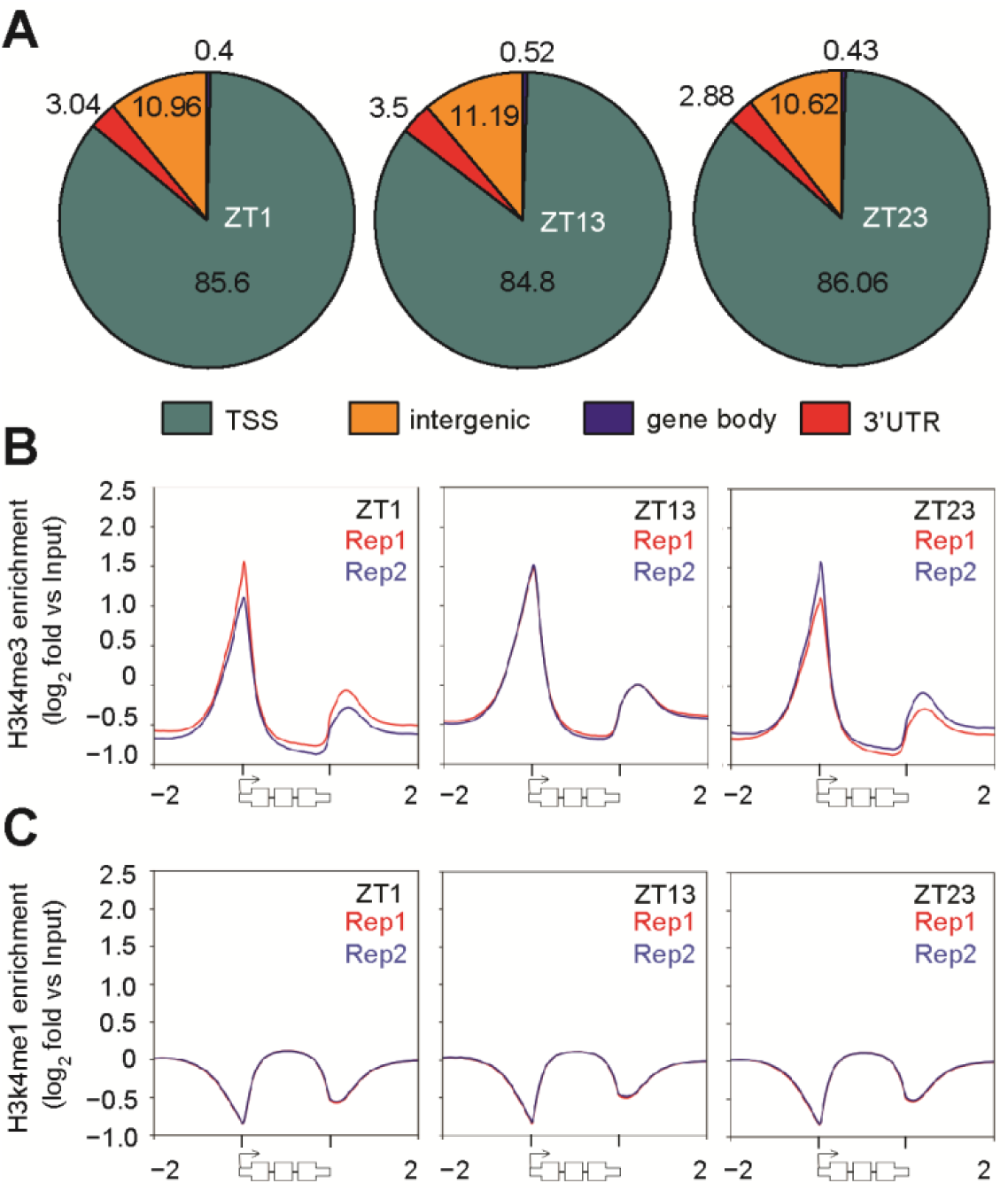
Genomic distribution of H3K4me3 and H3K4me1 in three time points during the cell cycle. **(A)** Enrichment of H3K4me3 at genomic regions at time points ZT1, ZT13 and ZT23 as indicated. A 750 bp window from the middle of each H3K4me3 peak was allowed to determine enrichment at genomic regions such as transcription start sites (TSSs, cyan), intergenic regions (orange), gene bodies (blue) and 3’UTRs (red). **(B,C)** Gene wide distribution of **(B)** H3K4me3 and **(C)** H3K4me1 along all Chlamydomonas genes was visualized using density blots. For each gene, the histone modification intensity is displayed along –2 kb to 2 kb regions around the TSSs. Shown are density blots of H3K4me3 enrichment from two independent replications in red or blue, respectively.

**Figure 3.**
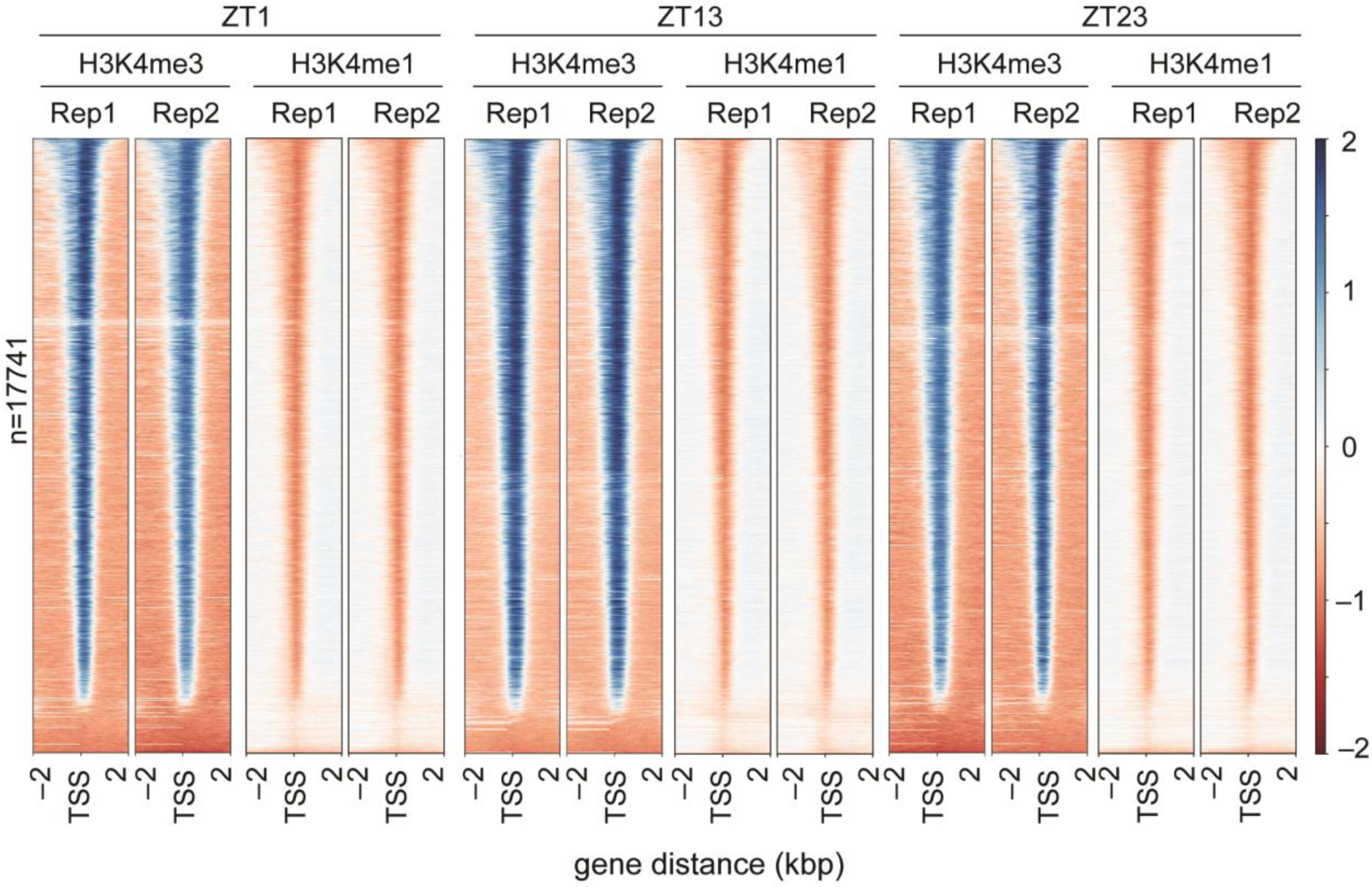
Distribution pattern of H3K4me3 and H3K4me1 around genes. The distribution of H3K4me3 and H3K4me1 along all Chlamydomonas genes in three time points during the cell cycle was visualized using heat maps. For each gene, the histone modification intensity is displayed along – 2 kb to 2 kb regions around the TSSs. Genes shown in both heat maps are sorted based on H3K4me3 enrichment. Blue color indicates enrichment over genomic DNA, red color indicates depletion as compared to genomic DNA. Rep = replicate.

### H3K4me3 is stably maintained independently of gene expression level for most genes

To investigate the relationship between gene expression and H3K4me3 TSS enrichment, we compared transcript abundances in FPKMs with H3K4me3 peak enrichment revealing a weak positive correlation between the two (R^2^ around 0.4, p-value ≤0.0001, Figure 4A). To further examine the nature of this correlation we divided all Chlamydomonas genes into six groups based on their transcript abundances, and then compared H3K4me3 enrichment for each group (Fig. 4B). This analysis showed that genes with very low expression (0-0.1 and 0.1-1 FPKM) were influencing the correlation, and that above an FPKM threshold of 1, there was almost no correlation between H3K4me3 and strength of gene expression over four orders of magnitude (1-10,000 FPKMs). A reciprocal analysis was also done where genes were grouped based on whether or not they had a detectable H3K4me3 peak at any of the three time points, and transcript abundance levels were compared between the two groups (Fig. 4C). This plot again showed that the small group of genes without any H3K4me3 peak at their TSS was significantly less expressed than those with a peak.

**Figure 4.**
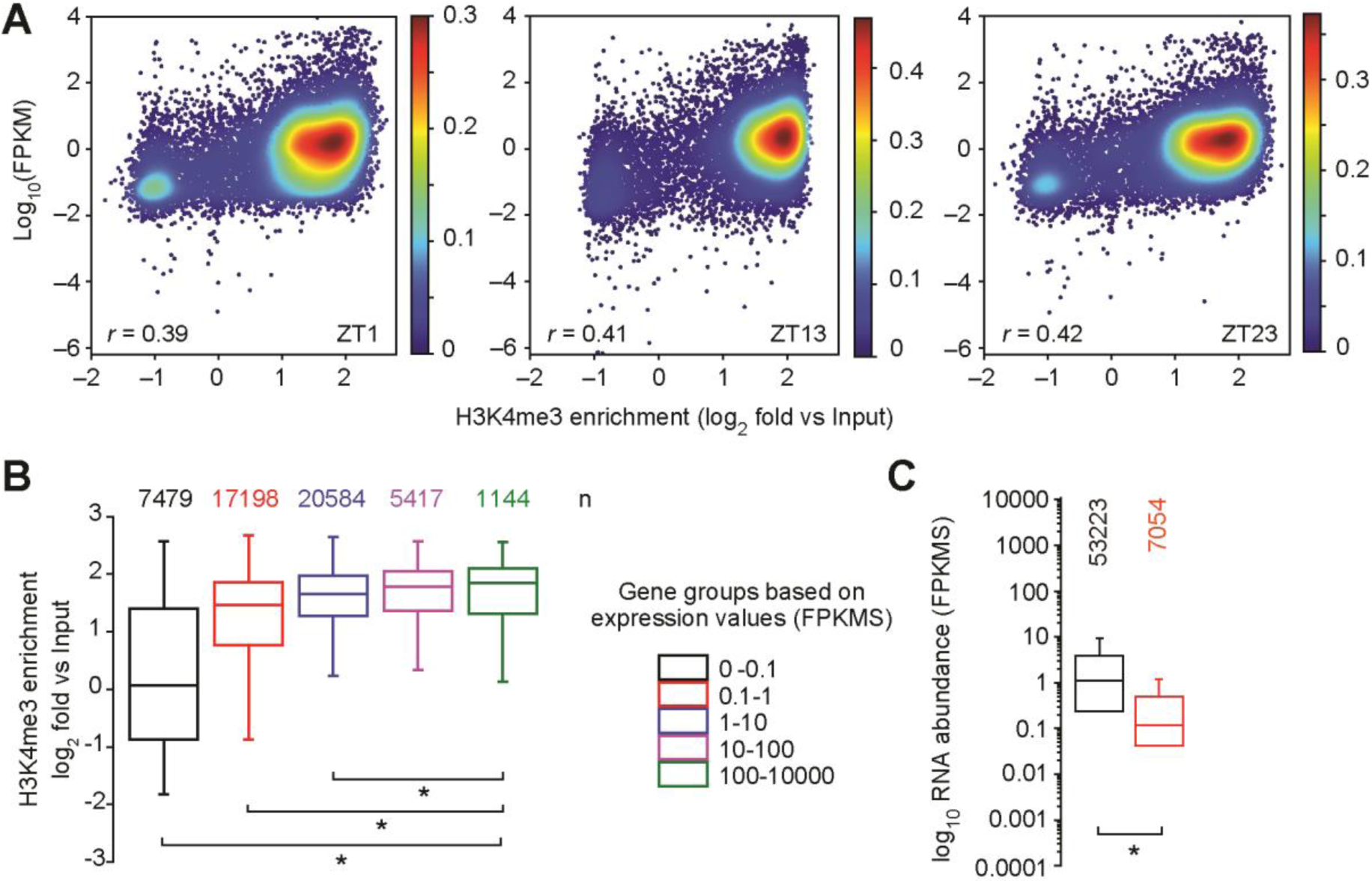
Relationship between H3K4me3 enrichment and gene expression. **(A)** Comparison of the ratio of transcript levels in FPKMs (determined by RNA-seq) of genes at all three time points with the ratio of H3K4me3 enrichment against input in the same time points. Shown is the Pearson correlation coefficient. **(B)** Levels of H3K4me3 enrichment in genes from five different groups of genes that were separated based on their respective transcript abundances in FPKMs. **(C)** Expression values in FPKMs of all genes across all three time points with H3K4me3 enrichment (black) and of genes that have no significant enrichment of H3K4me3 at their TSS (red). **(B,C)** *p < 0.05 (one-way ANOVA, Holm-Sidak post hoc test).

### Unexpected H3K4me3 enrichment patterns at infrequently expressed genes in the life cycle

The discovery that H3K4me3 peaks were largely uncoupled from gene expression magnitude led us to ask whether genes that were known to be repressed during the vegetative life cycle but well expressed under very specific conditions also retained H3K4me3 peaks. Sex is facultative in Chlamydomonas, and strains may replicate vegetatively for thousands of generations between matings. Several hundred early zygotic genes are expressed strongly and specifically in a short window after fertilization (35, 36), and are a good test for persistence of H3K4me3 peaks on genes that are very infrequently expressed (Supplemental table 2). Remarkably, nearly all these genes (83%) retained peaks despite very infrequent and unpredictable timing of expression (Fig. 5). Since zygotic genes were defined based on expression ratios compared with vegetative or gametic expression levels, some of them do have some basal vegetative or gametic expression. A more stringent filter for zygotic genes was created to retain only those genes that had low vegetative or gametic expression (<1 FPKM) and significant zygotic expression (>10 FPKM). In this list of 74 genes, 61 had H3K4me3 peaks at their promoters in at least one time point with 59 having a significant peak at all three time points (Supplemental table 2). Of the remaining 13 early zygotic genes, the absence of a detectable peak was due to missing sequence coverage at these genomic loci, so we found no evidence of a stringently-expressed early zygotic gene that lacked a H3K4me3 peak.

**Figure 5.**
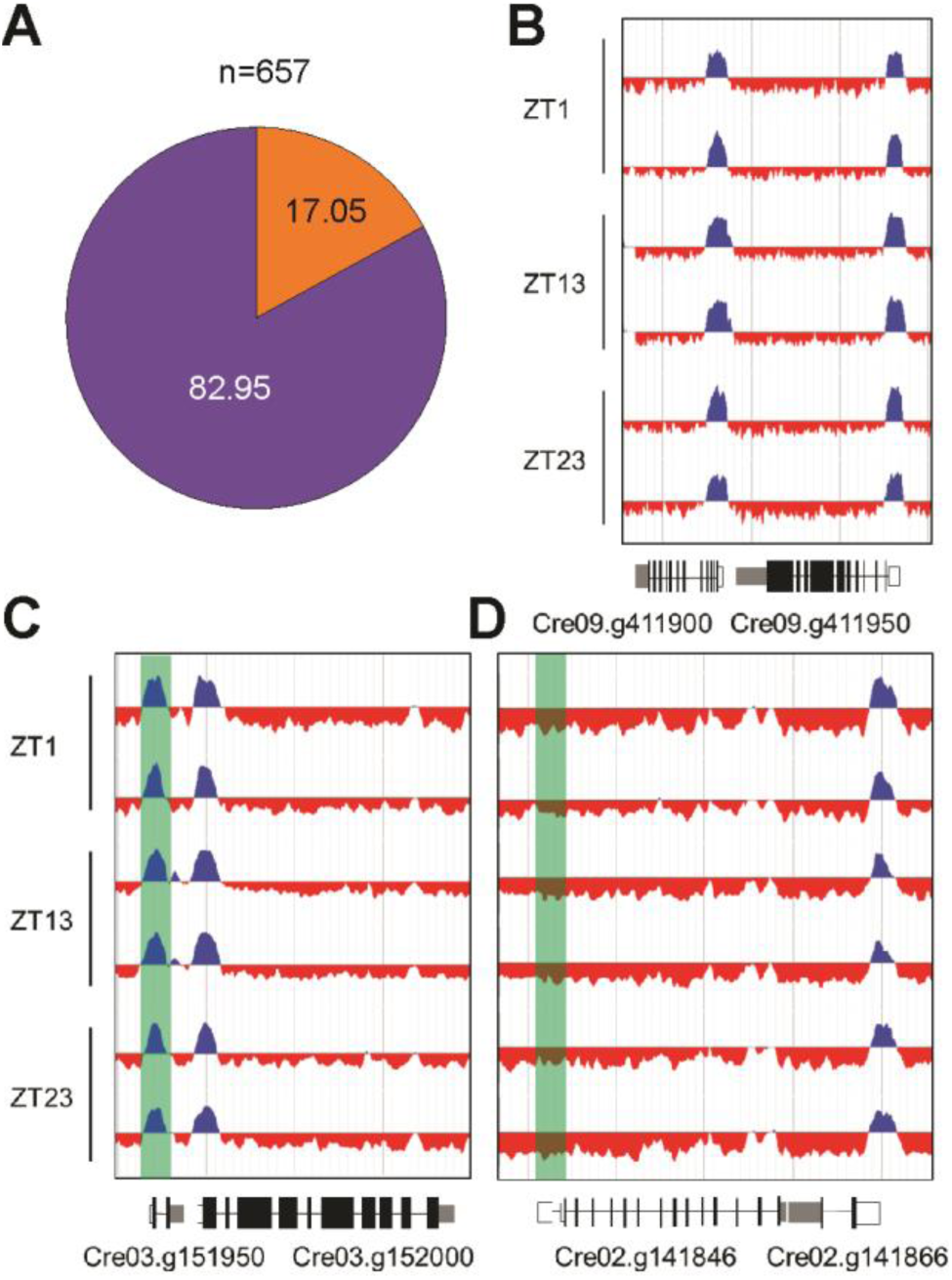
Presence and absence of H3K4me3 peaks at early zygotic and meiotic zygotic genes. **(A)** Proportions of 657 early zygotic genes that have a H3K4me3 peak at their TSS (purple) compared to those without H3K4me3 enrichment at their TSS (orange). **(B-D)** Replicate ChIP-seq tracks from samples taken at ZT1, ZT13 and ZT23 displaying H3K4me3 enrichment. Promoter regions of the respective genes are highlighted in green. Introns are drawn as black line, exons as black bars, 5’UTRs are shown as white bars, while 3’UTRs are shown as grey bars. **(B)** typical vegetatively expressed genes **(C)** a representative, early zygote gene and **(D)** a representative core meiotic gene encoding an ortholog of Rad51/DMC1 adjacent to a convergently transcribed vegetative gene that has a H3K4me3 peak. See Supplemental Table S2 for gene IDs and zygotic and meiotic gene expression data.

We also looked at H3K4 trimethylation on the TSS of nine well annotated core meiotic genes that are predicted to be expressed only when zygotes germinate. Seven of these meiotic genes— including those encoding single orthologs of Spo11, Hop1, Dmc1, and mismatch repair proteins— were completely missing H3K4me3 at their predicted TSS (Fig. 5). The two meiotic genes which retained H3K4me3 at their TSS, *HOP2/TBPIP* and *MND*, are also expressed in the mitotic cell cycle and are predicted to function in both meiosis and mitosis. It is not clear whether core meiotic genes have different epigenetic markings at their promoters, or whether they might transiently acquire H3K4me3 when expressed. It is also unclear whether other classes of genes are also missing this mark, but the absence of H3K4me3 at the TSS of well supported gene models whose expression is required during the life cycle could be a potential filter to identify new meiotic genes or other classes of genes that are functionally related to each other.

A third class of facultative genes are those expressed in response to abiotic stresses such as macro nutrient or trace element limitation. While there are many such genes documented in different studies (38-41), almost all of them had H3K4me3 peaks at their TSS, but also had significant basal expression even in the absence of a stress trigger, so their retention of H3K4me3 might not reflect epigenetic memory per se, but the presence of a basally active promoter at those loci.

In summary, the early zygotic genes we analyzed demonstrate the potential for long-term persistence of H3K4me3 at TSS of promoters over many generations even in the absence of expression, while the core meiotic genes are a noteworthy exception to the near ubiquitous presence of H3K4me3 at the TSS of nearly all genes, regardless of their expression status.

### Dynamic H3K4me3 peaks and their relationship to gene expression

While the vast majority of the H3K4me3 peaks that we identified were stably maintained over the course of the cell cycle, we found 120 genes with a dynamic pattern where peak intensity fluctuated in a cell-cycle-dependent manner (Supplemental table 3). We focused on 62 of these peaks that were located at TSS and found that the majority (45/62) had a strong H3K4me3 peak during cell division (S/M) phase with weak peaks during G1 and G0 (Fig. 6). Strikingly, genes with a dynamic S/M peak included 8 histone genes, a result that was unlikely to be obtained by chance among >17,000 predicted protein coding genes, even when factoring in that there are >30 histone gene clusters throughout the genome (Fig. 6). Core histone clusters in Chlamydomonas typically have bi-directional promoters, often with adjacent H2A/H2B and H3/H4 pairs, and are expressed in a strong burst during S/M phase and are termed replication-dependent (Fig. 7A, B)(33, 34). A small minority of histone loci, however, is replication independent and are expressed at constant low levels across the cell cycle (Fig. 7C) (33). Interestingly, the dynamic peak behavior we observed applied only to the replication dependent class of histone genes and not the replication independent ones (Fig. 7D-F and Supplemental Figure 1-4). Besides histone genes, we noted that several other genes were among those with dynamic S phase H3K4me3 peaks at their TSS and were also expressed most strongly during S/M, including the cell cycle gene *MCM10* (Fig. 6). However, having dynamic S/M phase H3K4me3 peaks was the exception for cell cycle genes and not the rule. An exceptional feature of replication dependent histone transcripts in Chlamydomonas is the absence of 3’ polyadenylation. To ask whether other non-polyadenylated transcripts also had dynamic H3K4me3 peaks, we surveyed H3K4me3 enrichment at genes coding for rRNAs, tRNAs and snRNAs. While rDNA arrays and tRNAs mapped to genomic regions with low read coverage and impeded our ability to draw robust conclusions, we could map all previously identified snRNAs (snRNA U1, U2, U4, U5 and U6) to the current Chlamydomonas genome version (43-45). To validate snRNA loci, we verified expression of each by manual curation on the genome browser using our mapped transcript data set (Supplemental table 4). Strikingly, with one exception, we could indeed identify dynamic S/M phase H3K4me3 enrichment at the 5 prime regions upstream of the snRNAs U1, U2, U4 and U5 (Supplemental Figure 4). The only exception was the 5 prime region of U4 snRNA, which was located so close to the promoter region of a neighboring gene that changes in H3K4me3 enrichment at the U4 promoter region could not be ascertained.

**Figure 6.**
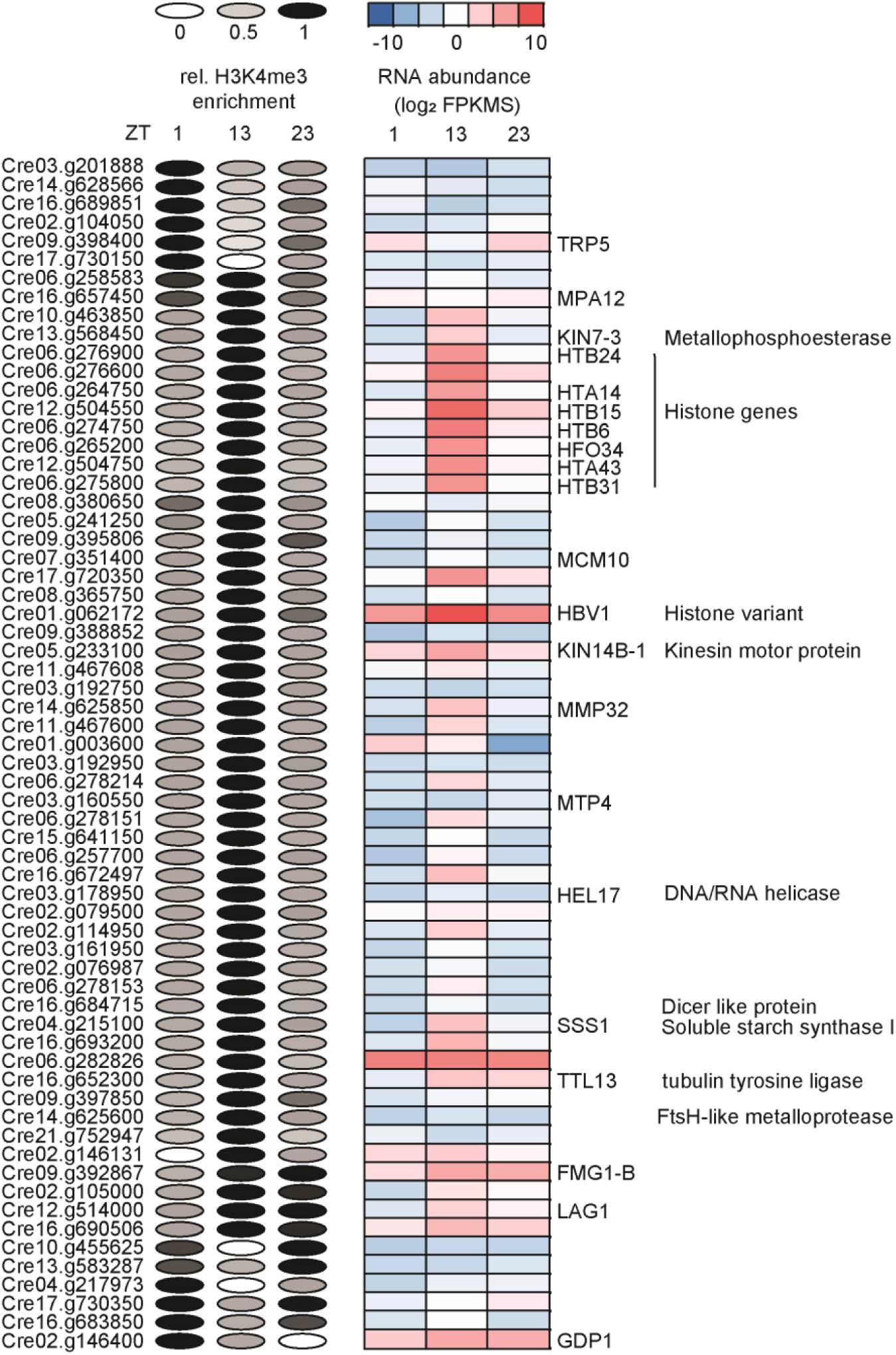
Overview of dynamic peaks spanning the TSS of genes. Left side: Relative H3K4me3 enrichment values at the TSS of the respective genes are visualized as ellipses. H3K4me3 enrichment was determined using EdgeR and values for replicates were averaged and then normalized to the time point with the maximum value that was set to 1. Relative peak height for each time point is represented by the gray scale value in each ellipse. Right side: RNA abundances of the respective genes are visualized as a heat map showing log_2_ FPKM values as averages of three replicates. Gene names and annotations were derived from Phytozome, (https://phytozome.jgi.doe.gov/pz/portal.html).

**Figure 7.**
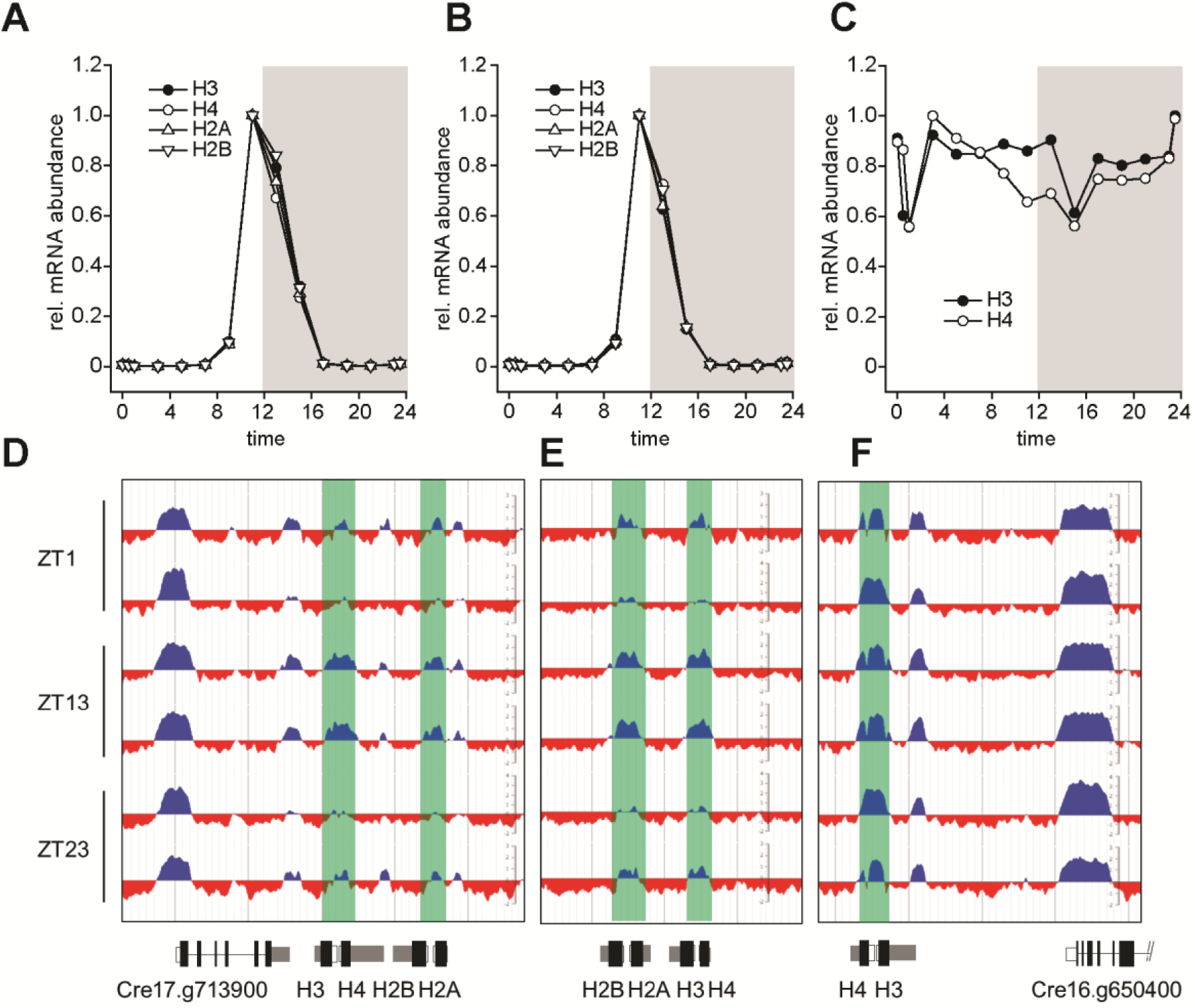
Cell cycle dependent expression and dynamic H3K4me3 peaks at core histone genes. Normalized expression patterns of **(A,B)** replication dependent histone clusters on chromosome 17 **(A)** H3 (Cre17.g713950), H4 (Cre17.g714000), H2A (Cre17.g714100), H2B (Cre17.g714050) and on chromosome 6 **(B)** H3 (Cre06.g268350), H4 (Cre06.g268400), H2A (Cre06.g268300), H2B (Cre06.g268250), as well as **(C)** replication independent constitutively expressed histones on chromosome 16, H3 (Cre16.g650300) and H4 (Cre16.g650250). (**D**) Relative expression values over the full diurnal cycle from ZT0 to ZT24 are based on averaging each of three replicates and setting the maximum expression value to 1 for each gene. **(D,E,F)** Replicate ChIP-seq tracks of histone loci in panels **A**-**C** with samples taken at ZT1, ZT13 and ZT23 as indicated. Introns are drawn as black line, exons as black bars, 5’UTRs are shown as white bars, while 3’UTRs are shown as grey bars.

In summary, we discovered a novel expression-associated dynamic H3K4me3 peak behavior for replication dependent core histone genes and snRNA loci that suggests the epigenetic environment and control mechanism for genes encoding non poly-adenylated transcripts may be different than that for typical protein coding genes.

### Relationship between histone H3 lysine 4 methylation and DNA methylation

In many species 5-methylcytosine (5meC) on nuclear DNA is associated with heterochromatin or gene silencing. Although Chlamydomonas has a much lower percentage of nuclear 5meC than other species where its function has been characterized (<0.75% of total C (36, 46), it has been localized to around two dozen hyper-methylated regions that have lower-than-average gene density and high repeat content, and which may correspond to heterochromatin (36). We analyzed the relationship between published regions of 5meC enrichment and H3K4 methylation status in our data. Intriguingly, we found that 5meC is enriched in regions of the nuclear genome that are lacking both H3K4me3 and H3K4me1 (Fig. 8A, B). The genes within these hypo-methylated regions are also enriched for those with poor expression (<1FPKM) compared with all genes (Figure 8C). In summary, our data point to an inverse correlation between H3K4 methylation and DNA methylation, suggesting that DNA hyper-methylated regions have atypical chromatin architecture.

**Figure 8.**
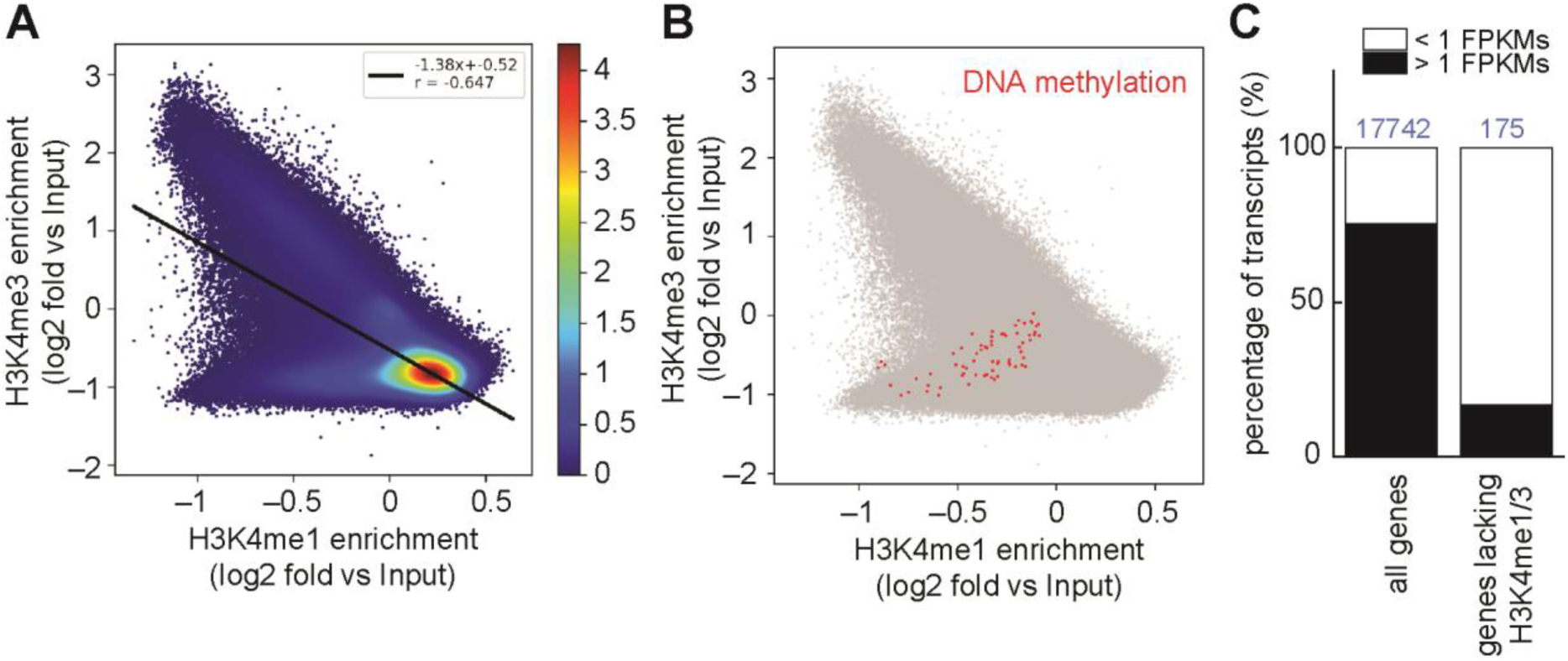
Correlations between H3K4 methylation, DNA cytosine-5 methylation and gene expression. **(A) Genome-wide** Comparison of H3K4me3 versus H3K4me1 enrichment plotted against input signal in (500 bp windows). The heat map scale depicts relative density of data points. **(B)** The same plot as in (A), but without the heat map scale and with 5meC-enriched loci plotted in red. **(C)** Stacked bar graphs represent the percentage of genes with high (>1 FPKM in at least one cell cycle time point) and low (<1 FPKM at all cell cycle time points) expression levels calculated for all 17742 predicted nuclear genes (left) and for the 145 genes located within hypermethylated genomic regions (right).

### H3K4me3 peaks as a genome annotation tool

Almost 11 percent of all identified H3K4me3 peaks mapped to intergenic regions in all three time points. We inferred that at least some of these peaks might be at unannotated loci.

Long non-coding RNAs (lncRNAs) are difficult to predict bioinformatically without empirical expression data and are easier to mispredict than protein coding genes since they lack long open reading frames. We focused on a large set of lncRNAs which were identified previously in Chlamydomonas (47) and were able to successfully map a total of 1067 to the current Chlamydomonas genome version 5.5 (Supplemental table 4). Around 40% of the predicted lncRNAs had a stable H3k4me3 peak at their TSS while 60% did not (Fig. 9A). Interestingly, the genes with H3K4me3 at their TSS were more likely to be expressed compared with those that did not, suggesting that presence of H3K4me3 is an effective filter for identifying the TSS and verifying expression of lncRNA loci (Fig. 9B).

**Figure 9.**
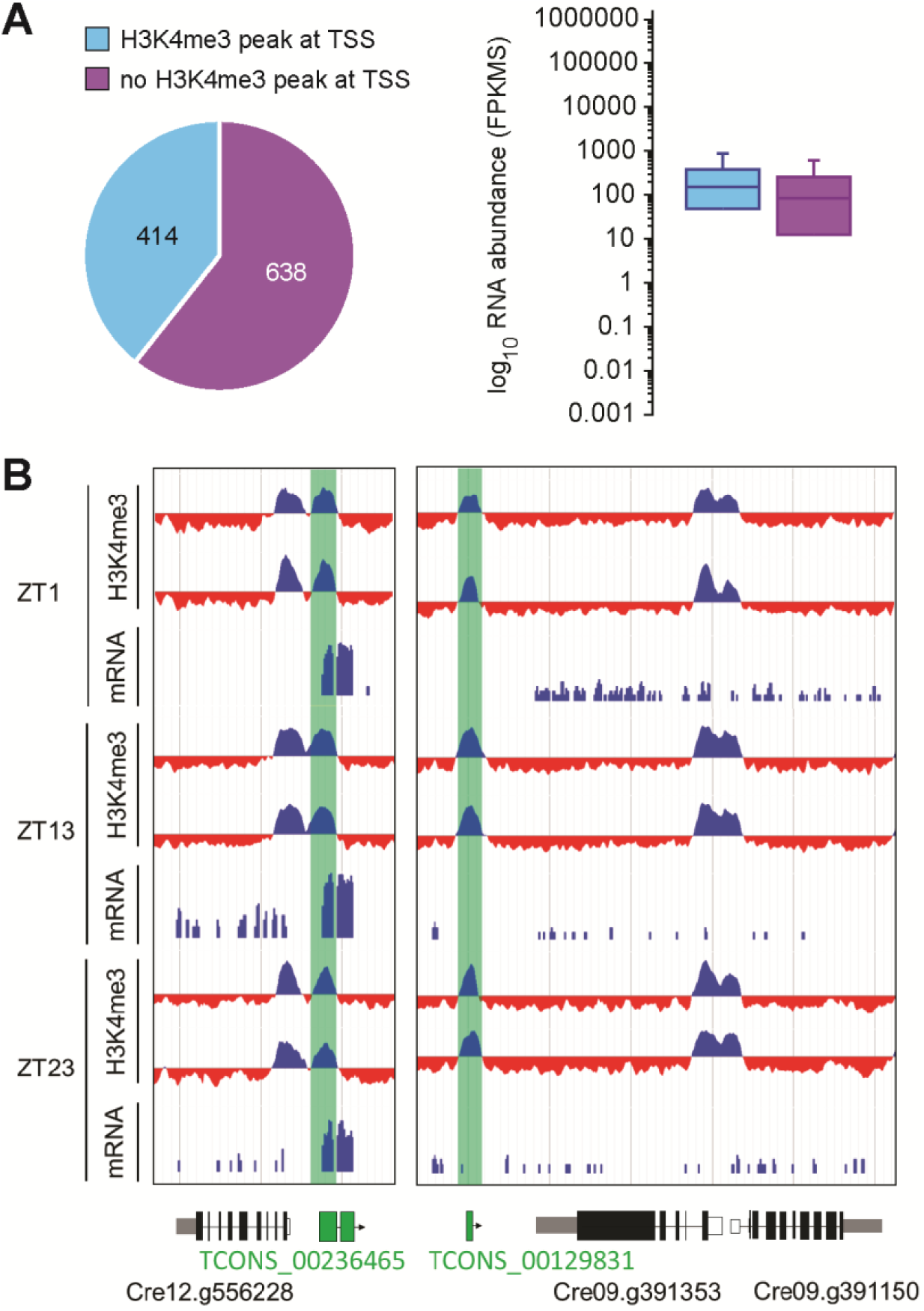
Validation of lncRNAs using H3K4me3 peaks at the TSS. **(A)** Pie chart showing the proportion of 1052 lncRNAs that have H3K4me3 enrichment at their TSS (turquoise) vs, without (purple) and boxplot visualizing RNA abundances in FPKMs of each group. **(B)** Replicate ChIP-seq tracks from samples taken at ZT1, ZT13 and ZT23 as indicated displaying H3K4me3 enrichment at TSSs of lncRNA (highlighted in turquoise). A clear H3K4me3 peak with no overlap to an annotated gene is highlighted in green. Introns are drawn as black line, exons as black bars, 5’UTRs are shown as white bars, while 3’UTRs are shown as grey bars.

## DISCUSSION

### Stable H3K4me3 enrichment at TSSs of the majority of genes during cell cycle progression

Our data demonstrates that H3K4me3 is a stably maintained chromatin mark in Chlamydomonas with enrichment at TSSs of the majority of known protein coding genes. This result further supports tri-methylated H3K4 enrichment at the transcription start sites of genes as a conserved feature in eukaryotes including yeast, mammals plants (8, 16, 18, 21, 48). In Chlamydomonas, H3K4me3 enrichment at TSSs correlates weakly with transcript abundances. More specifically, H3K4me3 seems to mark genes if their expression exceeded 0.1 FPKMs at least once during the cell cycle, even facultatively expressed genes, such as those involved in nutrient limitation responses and zygotic genes. Notably, genes that lack any expression evidence show a tendency to not be marked with H3K4me3 at their TSS. The correlation between H3K4me3 and gene expression may be related to expression potential (transcript levels when the gene is normally expressed) rather than frequency of expression. Extending this idea, many of the genes that lack H3K4me3 may not ever be expressed at high levels or have mis-predicted TSSs. Importantly, there are some genes that lack H3K4me3 but are almost certainly expressed in at least part of the life cycle. Core meiotic genes fall into this category and present an interesting exception. While H3K4me3 enrichment at TSSs of genes generally does not imply whether it is the cause or the result of underlying gene activation, we propose that H3K4me3 is one of the chromatin marks that, in combination with others, defines an active euchromatic state. Genes marked with H3K4me3 are poised for rapid activation, which allows for immediate transcriptional responses to changing environmental cues.

### Cell cycle dependent, dynamic H3K4me3 enrichment at histone gene promoters

While the majority of H3K4me3 peaks was maintained stably over the course of the diurnal cycle, less than one percent of all H3K4me3 peaks showed a more dynamic behavior. Notably, genes encoding histones are among the dominant protein coding gene class with a dynamic H3K4me3 peak at their TSS. The majority of Chlamydomonas histone gene clusters are expressed in a replication dependent manner during S/M phase. In Chlamydomonas and metazoans, replication-dependent histone mRNAs differ from canonical mRNAs in that they are not poly-adenylated, while by contrast, plants and many unicellular eukaryotes express only poly-A histone mRNAs (49-51). For a long time, the existence of non-polyA containing, replication-dependent expressed histone mRNAs was assumed to be a unique feature of metazoans until they were discovered in the green alga Chlamydomonas (51) and its relative Volvox (52). In Chlamydomonas, more than 29 copies of each of the core histone genes are organized in non-tandem gene clusters and expressed during replication (33, 51, 53). Strikingly, every single promoter for replication-dependent histone loci had H3K4me3 enrichment during mitosis, but the histone mark was absent during interphase. Interestingly, this observation is restricted to replication dependent expressed histone genes, as their constitutively expressed counterparts exhibited stable H3K4me3 enrichment at their TSSs during the cell cycle. We propose that dynamically marked histone gene promoters are depleted of H3K4me3 after mitosis by either the activity of histone demethylases or by histone turnover. Our observations suggest that histone gene promoters have some unique properties that may attract demethylases and/or induce active turnover of histones positioned near the TSS. How this phenomenon relates to histone gene expression remains to be determined.

### A unique, genome wide distribution pattern of H3K4me1 in Chlamydomonas

The genome wide distribution pattern of H3K4me1 seems to be unique in Chlamydomonas, in that almost the entire genome is mono-methylated at H3K4me1. While previous work based on LC-MS, and top down proteomics on Chlamydomonas histones indicated that mono-methylated H3K4me1 is a predominant histone PTM in Chlamydomonas (28-30), it is still striking to see the uniformity of this chromatin mark across the entire genome sparing only gene promoter regions. Notably, the genome wide distribution pattern of H3K4me1 seems to vary significantly between taxa, but the histone mark was mostly found within gene boundaries. For example in land plants like Arabidopsis, H3K4me1 was shown to cover entire gene bodies, but not the entire genome (21, 26). Our results in Chlamydomonas are in stark contrast to what has been observed in yeast, in which H3K4me1 is enriched at TSS (54). The same is true in humans, in which active genes are also characterized by high levels of H3K4me1 at their TSSs (16). Notably, H3K4me1 was also enriched at enhancers if H3K4me3 was depleted (20, 55).

### Inverse correlation between histone H3K4 methylation and DNA methylation

Most of the genome-wide DNA methylation and histone modification studies on mammalian cells show inverse correlation between DNA methylation and histone H3K4 methylation (56). Specifically, DNA methylation is associated with the complete absence of H3K4 mono-, di-, or tri-methylation (57). Our data shows a similar trend with an inverse correlation between the presence of H3K4 mono-and tri-methylation and DNA methylation. The latter indicates that DNA methyl transferases might be inhibited by H3K4 methylation also in Chlamydomonas, as this is the case in animal systems (58-62). The reciprocal relationship may also hold true, with methylated DNA regions being resistant to incorporation or retention of H3K4 methylated histones. While the mechanism(s) behind the antagonistic relationship of DNA methylation and histone methylation remains to be resolved, the ubiquitous presence of H3K4me1 may partly explain the paucity of DNA methylation in Chlamydomonas.

### High quality H3K4me3 map as an annotation tool

Our finding that the vast majority of protein coding genes have a discrete H3K4me3 peak at their TSS provides a potentially powerful annotation tool that can be used as positive verification for the start of transcription and for validation of predicted genes where other types of evidence might be lacking. In the case of lncRNAs, which cannot usually be verified based on similarity criteria, the presence of an H3K4me3 peak is also a potential verification tool. To date, the major challenge to identify novel lncRNAs in large transcriptome datasets is discriminating between possible transcriptional noise and physiological meaningful, dedicated expression (63, 64). It was noted that lncRNAs, like their protein coding counterparts, are associated with a specific chromatin domain which is characterized by the presence of trimethylated H3K4 within the promoter region and trimethylated H3K36 over the coding region, also referred as the so-called K4-K36 domain in mammals (65). Indeed, when we sought to functionally validate a catalogue of lncRNAs using H3K4me3 enrichment, we found that almost half of all candidate lncRNAs were associated with H3K4me3 at their TSSs. There seems to be no clear pattern as to why a lncRNA is associated with H3K4me3 enrichment or not. But, again like their protein coding counterparts, lncRNAs seem to be more likely to be trimethylated at H3K4me3 if they are actively transcribed. For either protein coding or non-protein coding genes, the absence of a H3K4me3 peak does not automatically discount a valid gene prediction. Conserved meiotic genes were missing this mark, and some genes such as those for canonical histones only had the mark at a specific time. Indeed, the absence of a H3K4me3 mark for genes that are otherwise predicted with high confidence may be highly informative about potential alternative expression mechanisms.

The relatively compact haploid genome (111 Mb) of Chlamydomonas combined with the ability to generate highly synchronous and uniform cell populations at different cell cycle and diurnal stages has previously enabled functional characterization of genes by co-expression analysis (66). Here we leveraged these highly advantageous properties Chlamydomonas to comprehensively characterize conserved H3K4 methylation marks across its genome at key cell cycle and diurnal time points, and compare them to DNA methylation patterns. Although most H3K4me3 peaks were stable, this synchronized system allowed us to identify a remarkable set of transient peaks at promoters of core histone genes. Overall, our findings highlight the potential for further discoveries coming from profiling additional epigenetic marks and histone modifications during the cell cycle.

## ACKNOWLEDGMENTS

Library preparation and sequencing (CSP1921), conducted by the U.S. Department of Energy Joint Genome Institute, a DOE Office of Science User Facility, and data analysis (DS) and manuscript preparation (DS) under the Laboratory Directed Research and Development Program of Lawrence Berkeley National Laboratory (SSM), are supported by the Office of Science of the U.S. Department of Energy under Contract No. DE-AC02-05CH11231. Sample preparation for JGI (DS) and bioinformatics at UCLA (AY) was supported by the Institute of Genomics and Proteomics under a cooperative agreement with the US Department of Energy Office of Science, Office of Biological and Environmental Research program under Award DE-FC02-02ER63421. Antibodies, project initiation / supervision, and manuscript editing was supported by the National Science Foundation MCB 1515220 and National Institutes of Health grant R01GM126557 to JGU.

## MATERIAL AND METHODS

### Chromatin-immunoprecipitation (ChIP) and library preparation

*Chlamydomonas reinhardtii* strain CC5390 was grown in a photo-bioreactor as described in (33). ChIP was performed as described in (67) with the following modifications: A total of 2×10^8^ cells, corresponding to 100 mL culture, were collected at the time points indicated by a 2-min centrifugation at 4°C and 3220g. The supernatant was discarded completely. To cross-link protein–DNA interactions, cells were resuspended in 10 mL freshly prepared cross-linking buffer (20 mM HEPES-KOH, pH 7.6, 80 mM KCl, and 0.35% formaldehyde) and incubated for 10 min at 24°C. Cross-linking was quenched by the addition of glycine to a final concentration of 125 mM and further incubation for 5 min at 24°C. Cells were collected by a 2-min centrifugation at 4°C and 3220*g*. Cells were lysed by the addition of 1000 μL lysis buffer (1% SDS, 10 mM EDTA, 50 mM Tris-HCl, pH 8.0, and 0.25× protease inhibitor cocktail [Roche]). Cells were sonicated on ice to generate an average DNA fragment size of ∼250 bp. ChIP was performed with aliquots corresponding to ∼2 × 10^7^ cells that were diluted 1/10 with ChIP buffer (1.1% Triton X-100, 1.2 mM EDTA, 167 mM NaCl, and 16.7 mM Tris-HCl, pH 8) and supplemented with 10 µg BSA. We omitted the addition of sonicated λ-DNA that was supplemented as blocking agent in (67) since it interferes with library preparation steps after ChIP. Antibodies specific for the following epitope were used: trimethylated H3K4 (10 μL, ab8580, lot# GR134854-1), monomethylated H3K4 (10 μL ab8895, lot# GR61306-1); Antibody-protein/DNA complexes were allowed to form during a 1-h incubation at 4°C, complexed with 6 mg pre-swollen protein A Sepharose beads (Sigma-Aldrich) during a 2-h incubation at 4°C, and precipitated by a 20-s centrifugation at 16,000*g*. Sepharose beads were washed once with washing solution 1 (0.1% SDS, 1% Triton X-100, and 2 mM EDTA, pH 8) containing 150 mM NaCl, once with washing solution 1 containing 500 mM NaCl, once with washing buffer 2 (250 mM LiCl, 1% Nonidet P-40, 1% Na-deoxycholate, 1 mM EDTA, and 10 mM Tris-HCl, pH 8), and twice with TE (1 mM EDTA and 10 mM Tris-HCl, pH 8). Protein-DNA complexes were eluted by incubating twice for 15 min at 65°C in elution buffer (1% SDS and 0.1 M NaHCO_3_), and cross-links were reverted by an overnight incubation at 65°C after addition of NaCl to a final concentration of 0.5 M. Proteins were digested by incubating for 1 h at 55°C after the addition of proteinase K (3.5 μg/mL), EDTA (8 mM), and Tris-HCl, pH 8.0 (32 mM). DNA was extracted once with phenol/chloroform/isoamyl alcohol (25:24:1), once with chloroform/isoamyl alcohol (24:1), and precipitated by incubation with 2 volumes of ethanol after addition of 0.3 M Na-acetate, pH 5.2, and 10 μg/mL glycogen over night at −20°C. Precipitated DNA was collected by a 20-min centrifugation at 4°C and 16,000*g*, washed with 70% ethanol, and air-dried and resuspended in 50 uL MilliQ water. 1-8 ng of ChIPed DNA was treated with end-repair, A-tailing, and ligation of Illumina compatible adapters (IDT, Inc) using the KAPA-Illumina library creation kit (KAPA biosystems). The ligated products were enriched with 8-10 cycles of PCR (HiFi premix, KAPA biosystems) and size selected to 200-500 bp with AMPure XP beads (Agencourt). qPCR was used to determine the concentration of the libraries. 5 libraries were pooled and sequenced on the Illumina Hiseq (HiSeq-1Tb at 1 × 100).

### ChIP-seq data analysis

The filtered reads from each library were mapped to the *Chlamydomonas reinhardtii* genome v5.5. H3K4me3 and H3K4me1 data were normalized against their corresponding genomic DNA inputs by using the BamCompare function in DeepTools (68). Average normalized signal density profiles over normalized lengths of all genes, as well as average normalized signal density profiles and heatmaps around TSS regions (± 2kb) of all genes (all together, or grouped into 10 deciles according to gene lengths) were generated by using the PlotProfile and PlotHeatmap functions in DeepTools (68). For the log_10_(FPKM) vs H3K4me3 normalized signal scatter plot, the mean normalized H3K4me3 signal value around TSS regions (± 250bp) for each gene was calculated and plotted against their log_10_(FPKM) values. For the H3K4me3 vs H3K4me1 normalized signals scatter plot, mean normalized signals over 1000 bp regions across the genome were calculated using the multiBigwigSummary function in DeepTools. The same function was also used to calculate the average normalized H3K4me1 and H3K4me3 signals over the previously reported hypermethylated regions (36).

H3K4me3 peaks were called using MACS2 (42), and H3K4me1 negative peaks were identified using in-house scripts. For the detection of negative H3K4me1 peaks, first the genomic bins with negative normalized signals were extracted and merged into continuous genomic regions using the bedtools merge function (69). Next, mean signal values and the minimum signal values over each region were calculated. Using these values, two z-scores were calculated for each region in comparison to genome-wide average; one for their mean signal, and one for their minimum signal. If a region’s either z-score was lower than -2, then that region was identified as the negative signal peak region. Regions that are shorter than 200 bp were discarded in this analysis. For both H3K4me3 and H3K4me1 peaks, the consensus peaks between two replicates at each time point were found using the bedtools intersect function (69). The same function was also used to identify peaks that overlap genes. In addition, H3K4me3 peak summit regions were checked against the gene regions (TSS, gene body, 3’UTR, and intergenic) using the bedtools closest function (69). If a peak summit was within a 750 bp window of any region, then the summit was mapped to that particular region.

Dynamic H3K4me3 peaks were identified using DiffBind. All the statistical routines available in the package were performed separately (EdgeR, DESeq, and DESeq2) between all pair-wise time points (ZT1 vs ZT13, ZT1 vs ZT23, ZT13 vs ZT23) and peaks with FDR values lower than 0.05 were identified as dynamic. All identified dynamic peaks were merged and further curated manually by visual inspection. Regions that are flagged as anomalous based on their extremely high (or extremely low) input read counts were discarded from the dynamic peak analysis. The anomalous regions were detected using the GreyListChIP package with 0.9 and 0.1 p-value thresholds for marking the regions as “grey” for their high and low read counts, respectively. Previously reported lncRNA sequences (47) were mapped to the soft-masked *Chlamydomonas reinhardtii* genome v5.5 using BLAT (70). 1085 out of 1440 reported lncRNA sequences were mapped with >99% match to the chromosome regions in the genome when the size of maximum gap between tiles in a clump was set to 3. The FPKM values and average mean normalized H3K4me3 signals around TSS (±250 bp) for 1052 lncRNA regions were calculated using Cuffdiff (71) and DeepTools (68), respectively. Sequences, transcript abundances and H3K4me3 enrichment at TSSs of all 1085 lncRNAs are listed in Supplemental table 4.

