## Supplemental Figures 1-4 for "Genome wide profiling of histone H3 lysine 4 methylation during the Chlamydomonas cell cycle reveals stable and dynamic properties of lysine 4 trimethylation at gene promoters and near ubiquitous lysine 4 monomethylation"

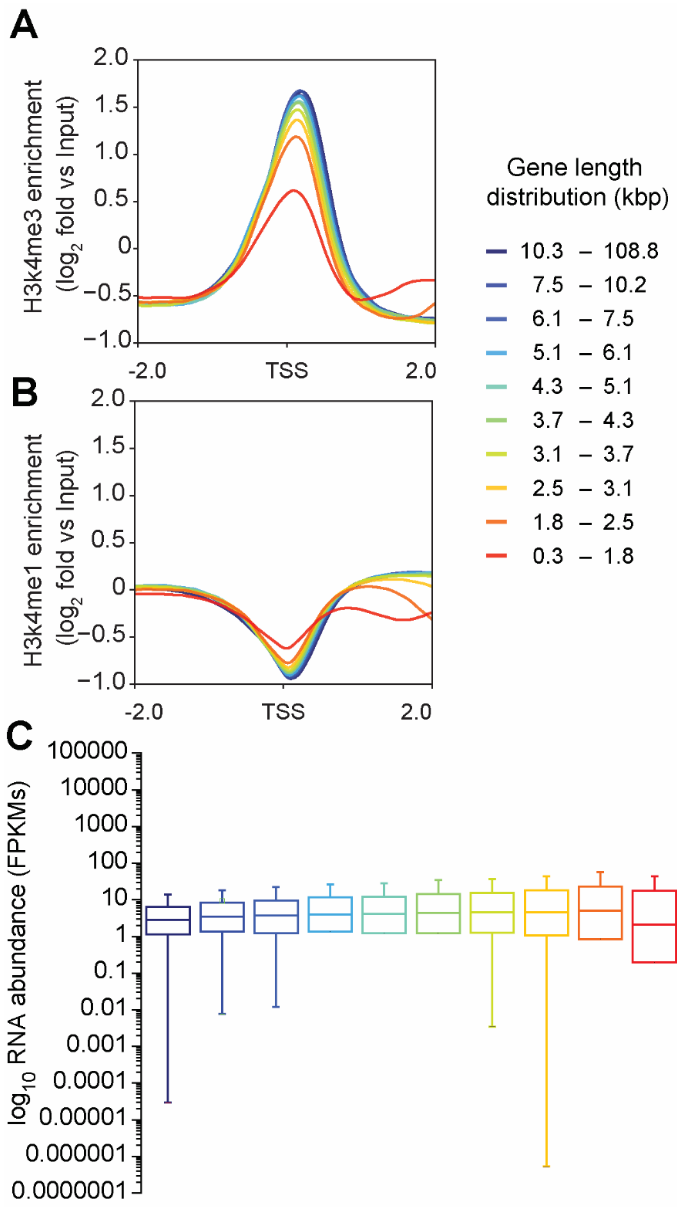


**Supplemental Figure 1. Correlation between gene length, histone lysine methylation and gene expression.** Density blots showing **(A)** H3K4me3 enrichment and **(B)** H3K4me1 enrichment against input in all time points. Genes were separated in groups by gene length from 1 to 10. **(C)** Boxplots showing expression values in FPKMS of all gene groups (1 to 10).

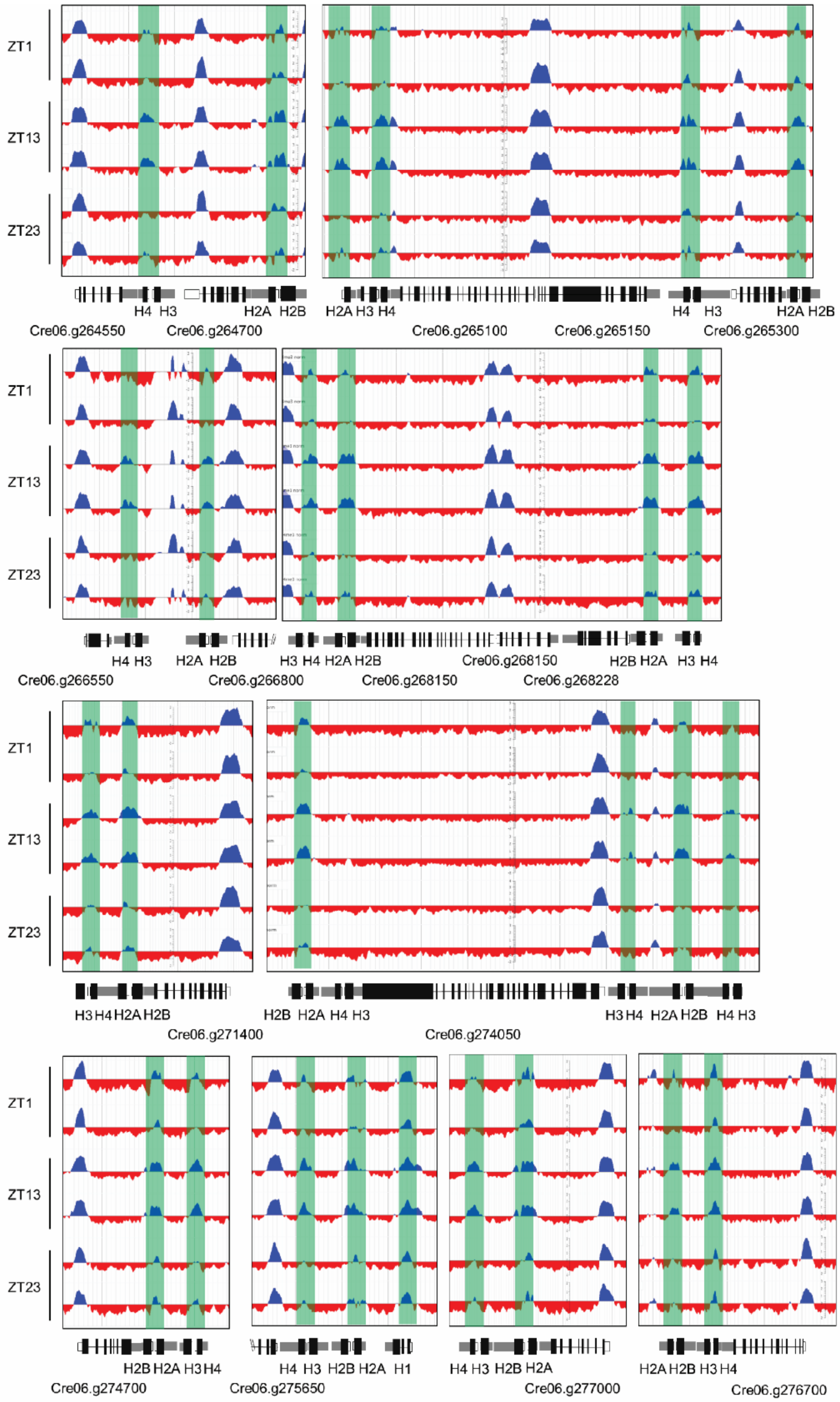


**Supplemental Figure 2. Genome browser view of dynamic histone H3K4me3 peaks at replication dependent expressed histone gene loci.** Shown are replicate ChIP-seq tracks from samples taken at ZT1, ZT13 and ZT23 as indicated. displaying H3K4me3 enrichment at histone gene promoters are highlighted in green. Introns are drawn as black line, exons as black bars, 5'UTRs are shown as white bars, while 3'UTRs are shown as grey bars.

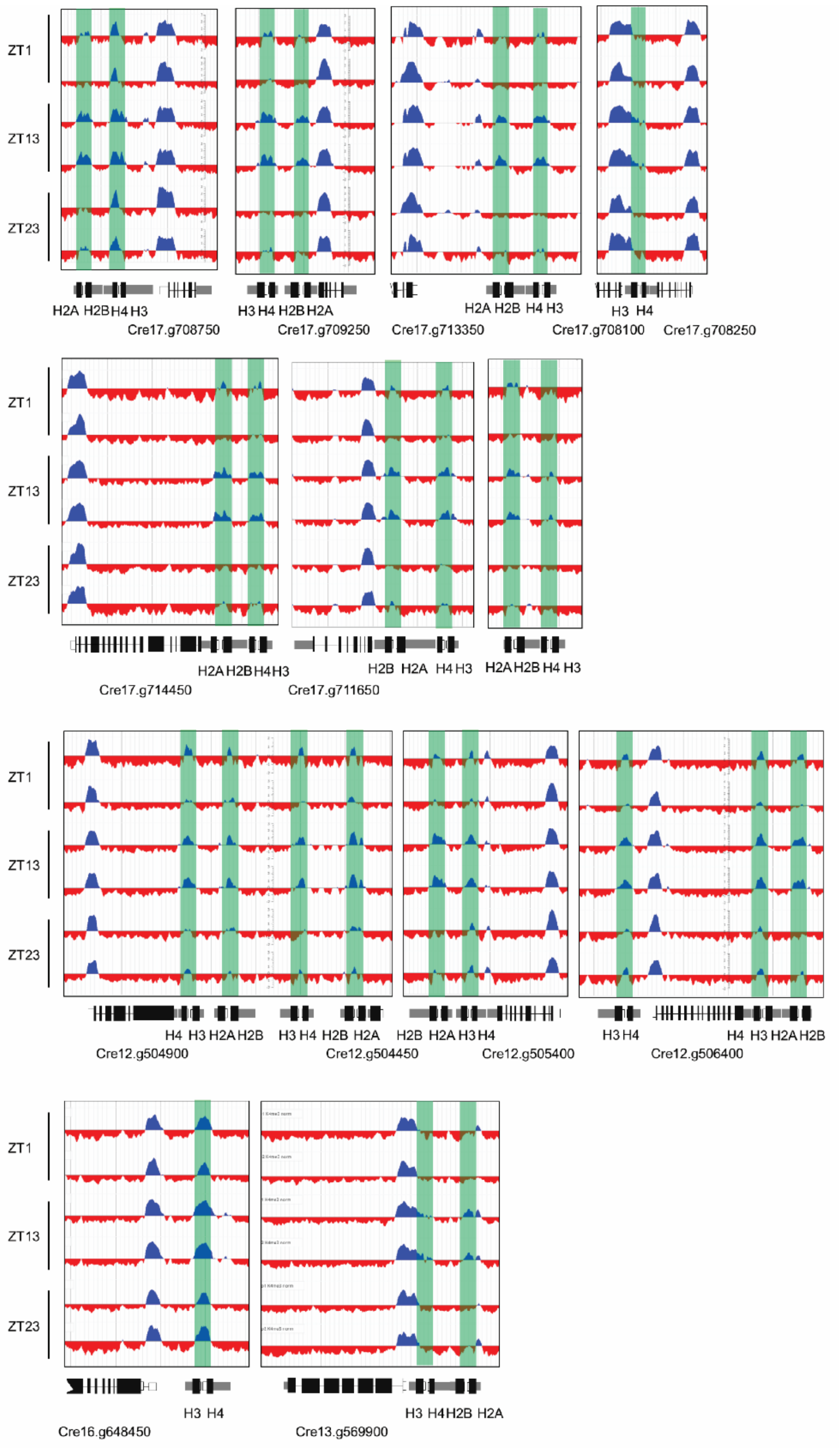


**Supplemental Figure 3. Genome browser view of dynamic histone H3K4me3 peaks at replication dependent expressed histone gene loci.** Shown are replicate ChIP-seq tracks from samples taken at ZT1, ZT13 and ZT23 as indicated. displaying H3K4me3 enrichment at histone gene promoters are highlighted in green. Introns are drawn as black line, exons as black bars, 5'UTRs are shown as white bars, while 3'UTRs are shown as grey bars.

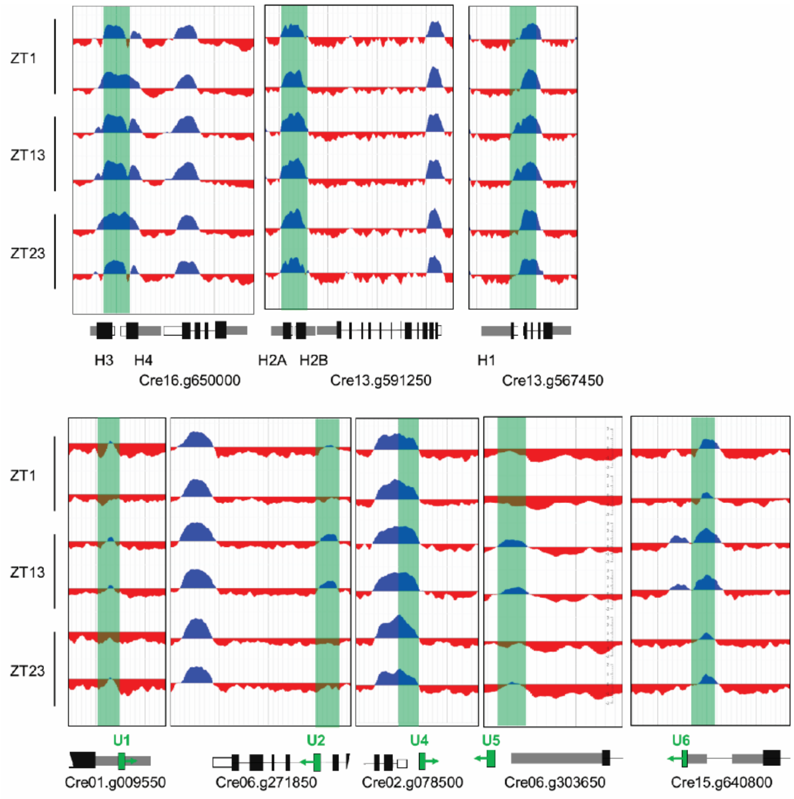


**Supplemental Figure 4.** Top panel: Genome browser view of stable histone H3K4me3 peaks at replication independent expressed histone gene loci**.** Bottom panel: Genome browser view of dynamic H3K4me3 peaks at promoter regions of snRNAs. Shown are replicate ChIP-seq tracks from samples taken at ZT1, ZT13 and ZT23 as indicated. displaying H3K4me3 enrichment at snRNA gene promoter regions highlighted in green. Introns are drawn as black line, exons as black bars, 5'UTRs are shown as white bars, while 3'UTRs are shown as grey bars.
